# Cyld aborts hyperactivation of synovial fibroblasts in inflammatory arthritis by regulating the TAK1/IKK2 signaling axis

**DOI:** 10.1101/2023.11.13.566552

**Authors:** Vagelis Rinotas, Kalliopi Iliaki, Lydia Pavlidi, Theodore Meletakos, George Mosialos, Marietta Armaka

## Abstract

TNF is a potent cytokine known for its involvement in physiology and pathology. In Rheumatoid Arthritis (RA), persistent TNF signals cause aberrant activation of synovial fibroblasts (SFs), the resident cells crucially involved in the inflammatory and destructive responses of the affected synovial membrane. However, the molecular switches that control the pathogenic activation of SFs remain poorly defined. Cyld is a major component of deubiquitination (DUB) machinery regulating the signaling responses towards survival/inflammation and programmed necrosis that induced by cytokines, growth factors and microbial products. Here we follow functional genetic approaches to understand how Cyld affects arthritogenic TNF signaling in SFs. We demonstrate that in spontaneous and induced RA models, SF-Cyld DUB deficiency deteriorates arthritic phenotypes due to increased levels of chemokines, adhesion receptors and bone-degrading enzymes generated by mutant SFs. Mechanistically, Cyld serves to restrict the TNF-induced hyperactivation of SFs by limiting Tak1-mediated signaling, and, therefore, leading to supervised NFkB and JNK activity. However, Cyld is not critically involved in the regulation of TNF-induced death of SFs. Our results identify SF-Cyld as a regulator of TNF-mediated arthritis and inform the signaling landscape underpinning the SF responses.

## Introduction

Rheumatoid Arthritis (RA) is a complex inflammatory disease irreversibly affecting synovial joints. The thin synovial membrane is getting inflamed, gradually reverting itself into a hyperplastic and invasive tissue mass namely pannus, which causes joint destruction. The synovial fibroblasts (SFs), a major mesenchymal cell type populating synovium, actively participate in the pannus by exhibiting a proinflammatory profile with migratory and tissue-invading properties leading the tissue destruction^1^.

We have previously shown that SFs exclusively receives the pathogenic load of TNF to orchestrate and arthritic disease TNFRI signaling in TNF-dependent modeled arthritis, such as *hTNFtg*, *TNF^ΔARE^* or Collagen Antibody Induced Arthritis (CAIA)^2,3^. Transcriptomic and epigenomic analysis of the *hTNFtg* SFs highlighted NFkB as a major component of inflammatory arthritic process^4^. In sheer concert with this finding, the SF-specific IKK2 targeting in the *hTNFtg* mouse uncoupled the function of IKK2 in both TNF-induced NFkB and death responses *in vivo*, highlighting the complexity in signaling events under chronic inflammatory conditions^2^. Interestingly, accumulating evidence points to the multiple molecular checkpoints that regulate the outcome of TNF signaling pathway either to survival/inflammatory signals through the activation of NFkB and MAPKs or cell death^5^, therefore regulating physiology and disease.

Ubiquitination is a posttranslational modification which tags ubiquitin via its C terminus to a target protein directing it either for degradation or becoming scaffolds, and consequently regulates signalling pathways. Ubiquitin chains is formed by constitutive attachment of ubiquitin to either of seven different lysine (K) residues (K6, K11, K27, K29, K33, K48, K63) or the N-terminal methionine (M1, Met1, linear chains)^6^. Remodelling of ubiquitin (Ub) loads of proteins is performed by specific ubiquitin ligases or deubiquitinases (DUBs). The most well-studied types of Ub chains are the K48, which predispose to proteosomal degradation, the K63, which affects activation status of proteins and the M1 which affects scaffolding properties of proteins^7^. The Ub-editing enzymes A20 (encoded by Tnfaip3), Otulin and Cyld are critical compoments with DUB activity modifying cytokine- and growth factor-induced signalling pathways^8^. Extensive studies causally linked mutations in A20 with several inflammatory conditions, including arthritic diseases^9^. More recently, it was exhibited that Otulin remove M1 ubiquitin chains to dictate signalling, and mutations in Otulin are consequently linked to an autoinflammatory syndrome in humans and lethal polyinflammatory consequences in mice^10–14^. Cylindromatosis (Cyld) has been originally described as a tumor suppressor gene mutated in familial cylindromatosis, predisposing to the development of cancerous lesions in skin^15–17^. The majority of detected CYLD mutations affect either the DUB activity or its expression levels^18^. Remarkably, genome-wide association studies additionally implicated CYLD polymorphisms in inflammatory diseases^19^. Functional genetic approaches revealed that mutated Cyld is associated not only to cancerous conditions, but also to immune homeostasis and inflammation^8,20^. By removing K63 and M1 ubiquitin chains Cyld regulates several signaling pathways^16,17,21^ acting downstream of TNF, TGFbeta, Wnt/β-catenin, Hippo, Notch signals such as c-Jun N-terminal kinase (JNK), p38 mitogen-activated protein kinase (p38 MAPK)), nuclear factor-кB (NFкB)^22–26^. Recently Cyld had been also exhibited to act as another checkpoint that prevent RIPK1-dependent programmed necrosis induced by TNF^27^.

Multiple studies demonstrated that the output of Cyld in very much stimuli- and cell-context specific. Even in the absence of mutations or polymorphisms, downregulation of Cyld has been detected in affected tissues of debilitating diseases such as NASH^28^ or inflammatory arthritis^29^. In arthritic synovial fibroblast, low Cyld levels are inversely correlated with the NFkB activity in RASFs^29^. However, mechanistic studies on the effects of SF-specific Cyld knockdown or loss of deubiquitinase activity *in vivo* remains underexplored. In this study, we pursued to decouple the Cyld involvement in the inflammatory behavior of SFs, by employing both acute and chronic TNF-dependent modeled arthritis coupled with genetic targeting and biochemical studies on deubiquinase-deficient Cyld SFs.

## Results

### TNF-mediated arthritis is deteriorated by mesenchymal Cyld deficiency

To study the role of *Cyld* in SFs we generated mice carrying a mesenchymal-specific homozygous deletion of *Cyld* exon 9 (designated *Cyld^M-Δ9^*, Cyld DUB deficient), by crossing mice containing floxed exon 9 of *Cyld*^30^ with *Col6a1-Cre* transgenic mice^3^. Exon 9 deletion results in the expression of a truncated form of *Cyld* which lacks deubiquitinating activity, but retains domains interacting with TRAF2 or NEMO^30^. The *Col6a1-Cre Cyld ^f/f^ (from herein referred as Cyld^M-Δ9/Δ9^*) mice were viable and fertile. The mice appeared normal in development with no apparent growth defects even at 12 months of age (not shown). To evaluate the mesenchymal-specific contribution of Cyld in regulating the TNF-modeled arthritis, we generated mice of the *hTNFTg Cyld^M-Δ9/Δ9^* genotype. The mice were viable and born in normal mendelian frequency. Interestingly, the mice growth was evidently retarded as early as 4 weeks of age, compared to non-Cre littermate controls (Fig 1a). The gross observation of the mice at the age of 6 weeks revealed excessive swelling and redness of ankle joints (Fig 1b) and deformity of wrinkle joints (Fig 1c).

**Figure 1.**
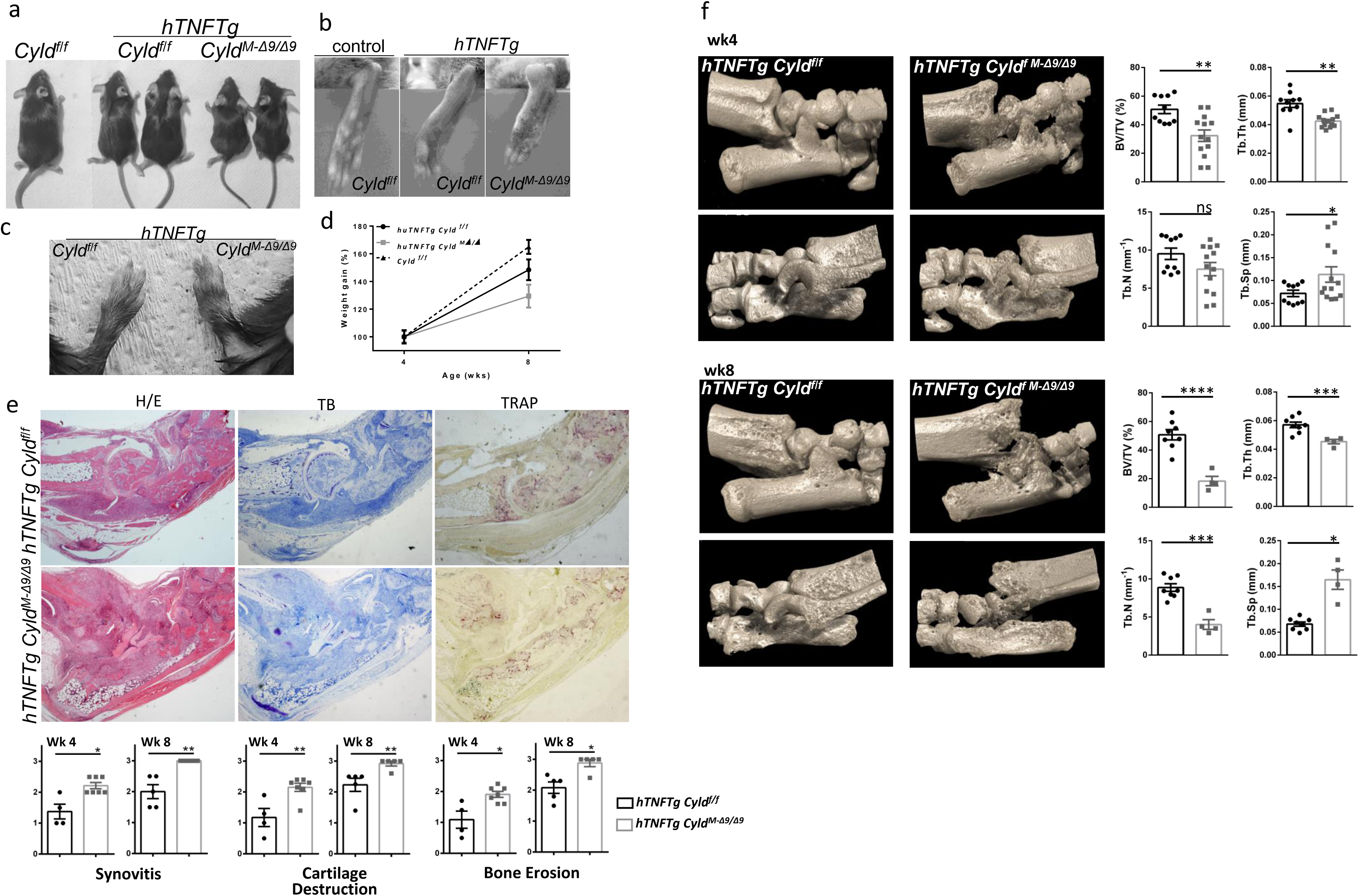
Mesenchymal Cyld deficiency deteriorates arthritic disease of *hTNFtg* mouse model. (a-c) Macroscopic view of Cyld^f/f^ and *hTNFtg Cyld^f/f^* and *hTNFtg Cyld^M-Δ9/Δ9^* mice at 6 weeks of age (a), their hind paws (b) and their forepaws (c). (d) Weight differences of Cyld^f/f^ and *hTNFtg Cyld^f/f^* and *hTNFtg Cyld^M-Δ9/Δ9^* mice at age of 4 and 8 weeks. (e) Histological examination of joints to evaluate inflammation (H/E panel), cartilage degradation (Toluidine blue-T/B panel) and bone erosions (Tartrate-resistant acid phosphatase-TRAP panel) in serial sections of 8-week old mice (n=5-7 mice/group). (f) mCT-based imaging and histomorhometric indexes for *hTNFtg Cyld^f/f^* and *hTNFtg Cyld^M-Δ9/Δ9^* mice at age of 4 and 8 weeks (n=9-13, week4; n=4-8, week 8). (BV/TV:Bone Volume Fraction, Tr. Th.: Trabecular thickness, Tb. N: Trabecular number, Tr. Sp: Trabecular Separation). Data are presented as the mean ± SEM. *P < 0.05, **P < 0.01 and ***P < 0.001 by two-tailed Student’s t-test (e-f).

To further evaluate the effects of mesenchymal-specific *Cyld* inhibition on arthritic features, histological sections of ankle joints from all groups of mice were analyzed in early (week 4) and intermediate phase of *hTNFtg* disease (8 weeks). Four-week old control *hTNFTg Cyld^f/f^* mice showed moderate inflammatory synovitis and mild cartilage degradation as it has been already reported for the original phenotype of *hTNFTg* mice^31^. In sharp contrast, the *hTNFTg Cyld^M-Δ9/Δ9^* mice showed severe inflammation of the synovium accompanied by marked proteoglycan loss of articular cartilage (Fig 1e). In later stages (8 weeks), the *hTNFTg Cyld^f/f^* mice develops full-blown manifestations of synovitis, cartilage destruction and bone erosions (Fig. 1e). At the same age, all the histological features of the ankle joints of *hTNFTg Cyld^M-Δ9/Δ9^* mice advocated for an extremely severe arthritic phenotype as this exhibited by the complete loss of joint architecture (Fig. 1e). The histomorphometric analysis fully supported the histopathological findings, with all bone parameters being heavily affected (Fig 1f).

To further confirm the role of mesenchymal Cyld-DUB deficiency in the development of arthritic phenotypes, we subjected *Cyld^M-Δ9/Δ9^*and littermate *Cyld^f/f^* control mice to (collagen antibody induced, CAIA)^2^ and we followed up the disease for 10 days (Fig. 2a). At both peak of disease (day 8) and the remission phase (day 10), we identified a deterioration of arthritic phenotype both macroscopically and histologically at the peak and disease and a delayed remission phase compared to Cyld sufficient littermate controls (Fig 2b). Collectively, these data clearly show that mesenchymal *Cyld* deficiency deteriorates arthritic manifestations in both innate- and autoimmune-based arthritis.

**Figure 2.**
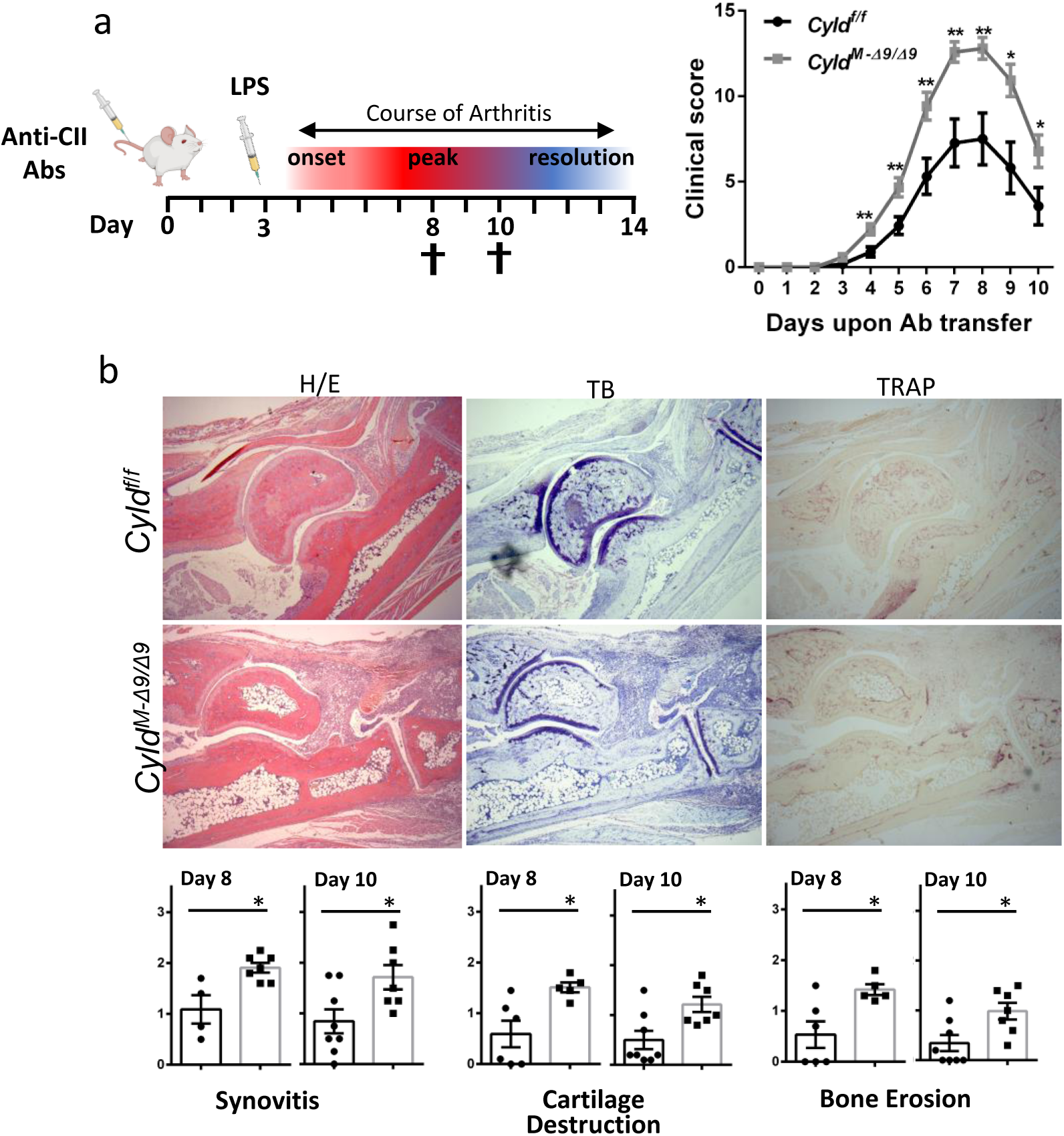
Mesenchymal Cyld deficiency exacerbates collagen-antibody induced arthritis (CAIA). (a) CAIA was induced in mice by injection of an anti-collagen antibody cocktail on day 0, followed by a single injection of 75 µg of LPS on day 3 (left panel). Semi-quantitative arthritis scores of of *Cyld^f/f^* and *Cyld^M-Δ9/Δ9^* mice upon CAIA induction (right panel) (n=14, two experiments) (b) Histological examination of joints to evaluate inflammation (H/E panel), cartilage degradation (Toluidine blue-T/B panel) and bone erosions (Tartrate-resistant acid phosphatase-TRAP panel) in serial sections of mice (n=7-8 mice/group). Photos of ankle joints depicted from Day 8 upon CAIA induction. Data are presented as the mean ± SEM. *P < 0.05, **P < 0.01 and ***P < 0.001 by two-tailed Student’s t-test.

### Cyld deficiency predisposes SFs to generate an amplified inflammatory signature

To better understand the effects of Cyld-DUB deficiency in the deterioration of arthritis, we analyzed major cellular composites characterizing arthritic joints during the early phase of *hTNFTg* arthritis (4 weeks of age). The relative abundance of lining (Thy1-) and sublining (Thy1+) synovial fibroblast types did not differ between the *hTNFTg Cyld^M-Δ9/Δ9^* and *hTNFTg Cyld^f/f^* genotypes, however both genotypes showed statistical differences compared to WT SFs (Fig 3a-left panel). Neutrophils and Ly-6C^low^ monocytes highly infiltrated the joints in early phases of arthritic disease and correlate with disease activity^32^. Interestingly, we detected higher number of both myeloid types in the *hTNFTg Cyld^M-Δ9/Δ9^ joints* compared to *hTNFTg Cyld^f/f^* controls (Fig 3a-right panel). These results suggest that the enhanced immune infiltrations in *hTNFTg Cyld^M-Δ9/Δ9^* joints rather than over-representation of SFs, characterize the accelerated disease onset.

**Figure 3.**
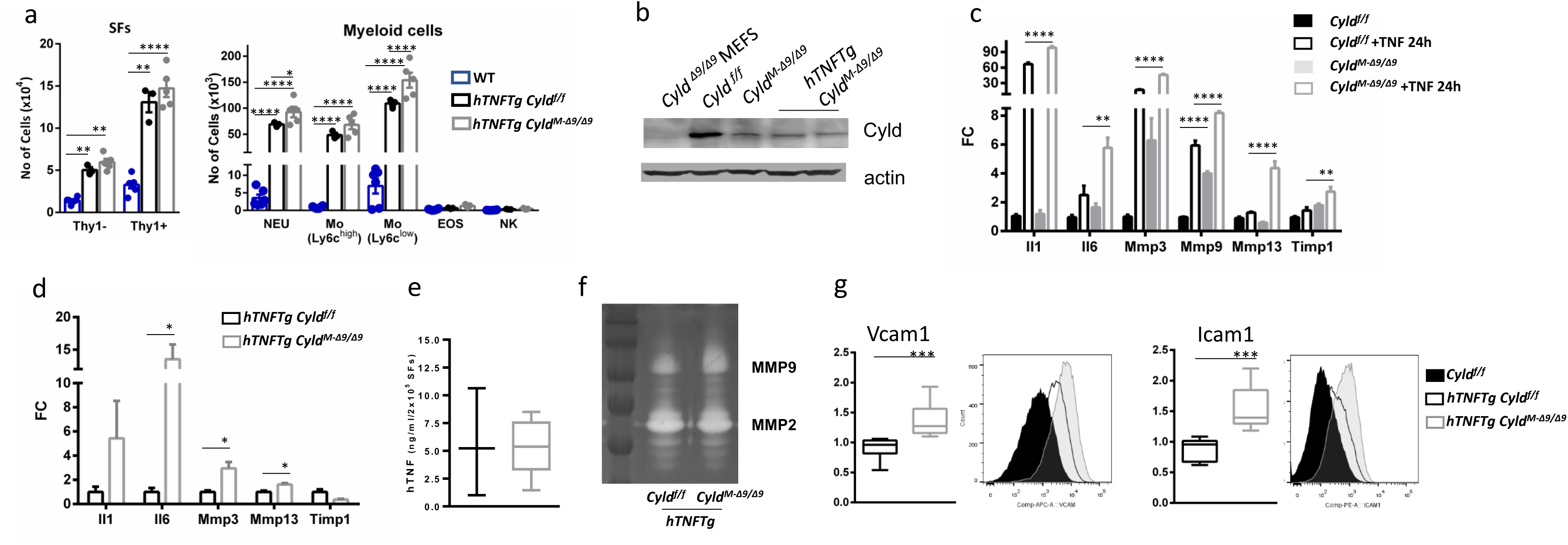
Cyld-DUB deficient SFs exhibit enhanced expression of arthritic mediators. (a) Flow cytometric analysis of SF subtypes (Τhy1-, Lining SFs; Thy1+, Sublining SFs) and myeloid infiltrates in the ankle joints of *hTNFtg Cyld^f/f^* and *hTNFtg Cyld^M-Δ9/Δ9^*4 week-old mice compared to WT controls. (b) Cyld expression in SF cultures derived from indicated genotypes. Extracts from Cyld-deficient MEFs served as control. (c) Relative gene expression of arthritic mediators *Il1b, Il6*, *Mmp-3*, *Mmp9, Mmp13*, *-Timp1* (normalized to b2m levels) in SFs of Cyld-proficient and Cyld DUB-deficient SFs upon TNF (20 ng/ml, 24 h) (*n* = 4) (d) Relative gene expression of arthritic mediators *Il1b, Il6, Mmp3, Mmp13 and Timp1* (normalized to b2m levels) in SFs of *hTNFtg* Cyld-proficient and Cyld DUB-deficient SFs. (e) hTNF protein levels in cultures of *hTNFtg* Cyld-proficient and Cyld DUB-deficient SFs (n=3-6). (f) Representative gelatin-based zymography employing supernatants of SF cultures derived from *hTNFtg* Cyld-proficient and Cyld DUB-deficient SFs. (g) Flow cytometric detection of Vcam-1 and Icam-1 levels in cultured *hTNFtg Cyld^f/f^* (black), *hTNFtg Cyld^M-Δ9/Δ9^* (grey) (upper panel) and representative FACS plots (*n* = 8, 2 experiments). Data are presented as the mean ± SEM (or SD for e) *P < 0.05, **P < 0.01, ***P < 0.001 and ****P<0.0001 by two-way ANOVA (Bonferroni correction) (a) one-way ANOVA (Bonferroni correction) (c) or unpaired t test with Welch’s correction (d and g).

To determine how Cyld-DUB deficient SFs affect the recruitment and the tissue destruction in the arthritic joints, we examined SF responses *ex vivo*. Upon confirming the loss of *Cyld* signals in cultured SFs isolated from ankle joints of mice of both genotypes (Cre and non-Cre littermates) at the age of 6-8 weeks (Fig 3b), we analyzed gene expression of known regulators of arthritic disease. TNF stimulated Cyld-DUB deficient SFs were highly responsive to TNF signals compared to the Cyld sufficient SFs, expressing higher levels of cytokines and proteases (Fig 3c). Interestingly, the levels of Mmp9 mRNA in Cyld-DUB deficient SFs were higher even in naïve conditions. Similarly, *hTNFTg Cyld*-deficient cells showed a particularly high expression of IL-6 as well as disbalanced expression ratio of the proteases Mmp3, Mmp13 and Timp1 (Fig 3d). IL1 was not significantly altered, but steadily tended to show higher expression. Moreover, supernatants of the *hTNFTg Cyld*-deficient SF cultures showed higher MMP9 activity when analyzed by zymograms (Fig 3e). Remarkably, the expression levels of hTNF transgene remained unaltered (Fig 3d). Expression of adhesion molecules VCAM-1 and ICAM-1 were similarly elevated, when compared to *hTNFTg Cyld* sufficient cells (Fig 3f). These results suggest that Cyld-DUB regulates gene expression upon inflammatory stimuli in SFs and correlate well with the exacerbated arthritic phenotype of *hTNFTg Cyld^M-Δ9/Δ9^*mice.

### Cyld deficiency causes aberrant TNF-mediated signaling responses in SFs

We analyzed signaling events that underline the arthritic gene expression. As previously reported for Cyld-DUB deficient MEFs, the Cyld DUB-null SFs exhibited apparent aberration in NFkB and JNK responses upon TNF stimulation, characterized by enhanced and prolong phosphorylation of Jnk1/2 kinases, as well as Ikk2 kinase followed by altered IkB degradation pattern. Interestingly, ERK and p38 activation was less affected (fig 4a). Consistently, the nuclear extracts from *Cyld* deficient SFs exhibited increased DNA binding activity to NFkB binding sites compared to Cyld proficient SFs (Fig 4b). We also confirmed the aberrant activation of JNK and NFkB in arthritic Cyld DUB-null SFs (Fig 4c). Collectively, the above results demonstrate that, upon TNF stimulation, Cyld DUB inactivation in SFs, causes enhanced NFkB and JNK responses, suggesting that Cyld deficiency renders SFs hypersensitive to TNF-mediated signals.

**Figure 4.**
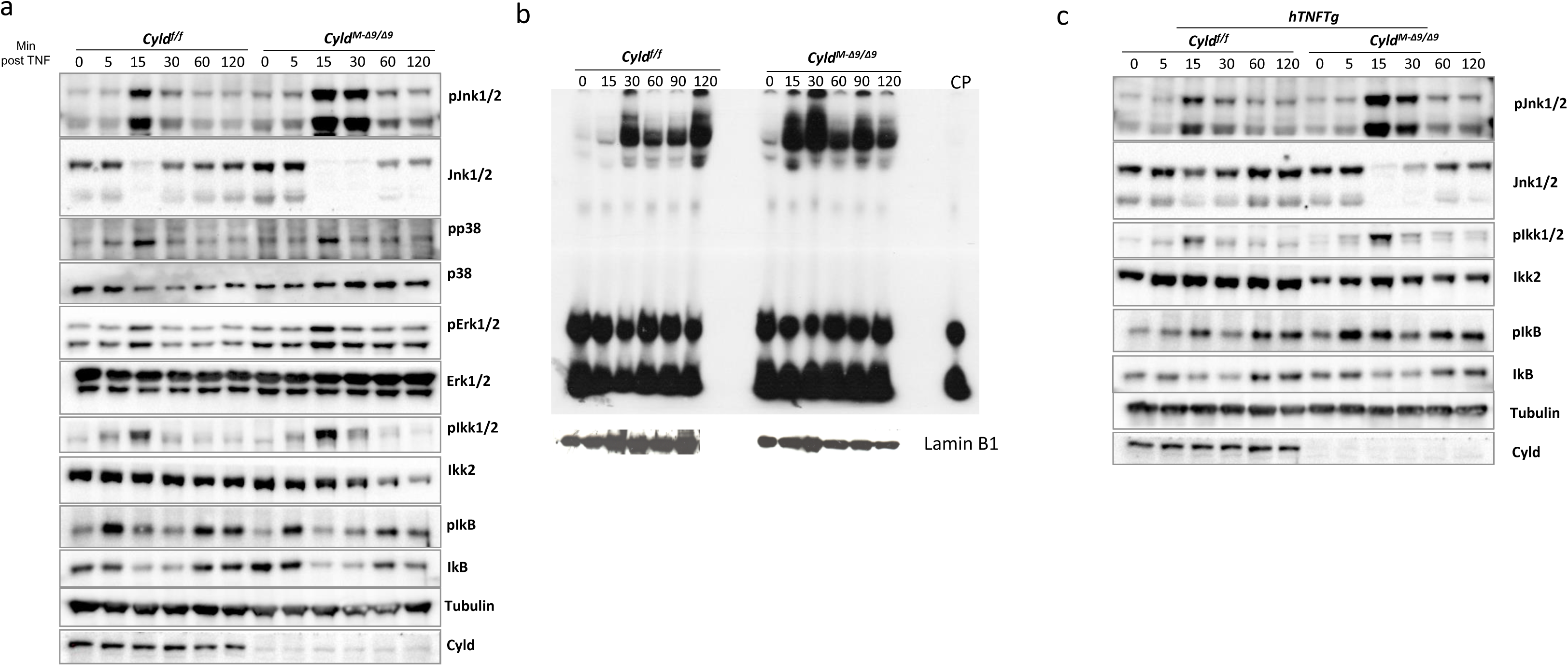
Cyld-DUB deficient SFs exhibit alter TNF-mediated signalling outputs. (a) Western blot analysis for the detection of MAPK/SAPK and NFkB activation in *Cyld^f/f^* and *Cyld^M-Δ9/Δ9^* cultured SFs upon TNF stimulation (10ng/ml) for the indicated time points. (n=5) (b) Band shift assay for the detection of NFkB binding activity of nuclear extracts derived from *Cyld^f/f^* and *CyldM-Δ^9/Δ9^* SF cultures, upon TNF stimulation for the indicated time points (n = 3). The same amount of extracts were subjected to Western blot analysis of LaminB1 expression. (CP: cold probe). (c) Western blot analysis for the detection of JNK and NFkB activation in *hTNFtg Cyld^f/f^* and *hTNFtg CyldM-Δ^9/Δ9^* cultured SFs upon TNF stimulation (10ng/ml) for the indicated time points. (n=5)

### Mesenchymal Cyld targets TAK1/IKK2 axis to regulate proinflammatory SF responses in TNF-mediated arthritis

To determine the molecular basis of aberrant TNF-mediated NFkB and JNK responses due to Cyld deficiency, we sought to examine the ubiquitinated state of TAK1, a kinase with shared regulatory features within these two pathways and a target of Cyld. In the absence of stimuli, TAK1 remained non-ubiquitinated and non-phosphorylated in both control and Cyld targeted SFs. Tak1 ubiquitination and phosphorylation peaked within seven minutes following stimulation in Cyld-proficient SFs and reduced to minimal within thirty minutes, however, we have observed sustained Tak1 phosphorylation in Cyld-DUB deficient SFs (Fig 5a, b).

**Figure 5.**
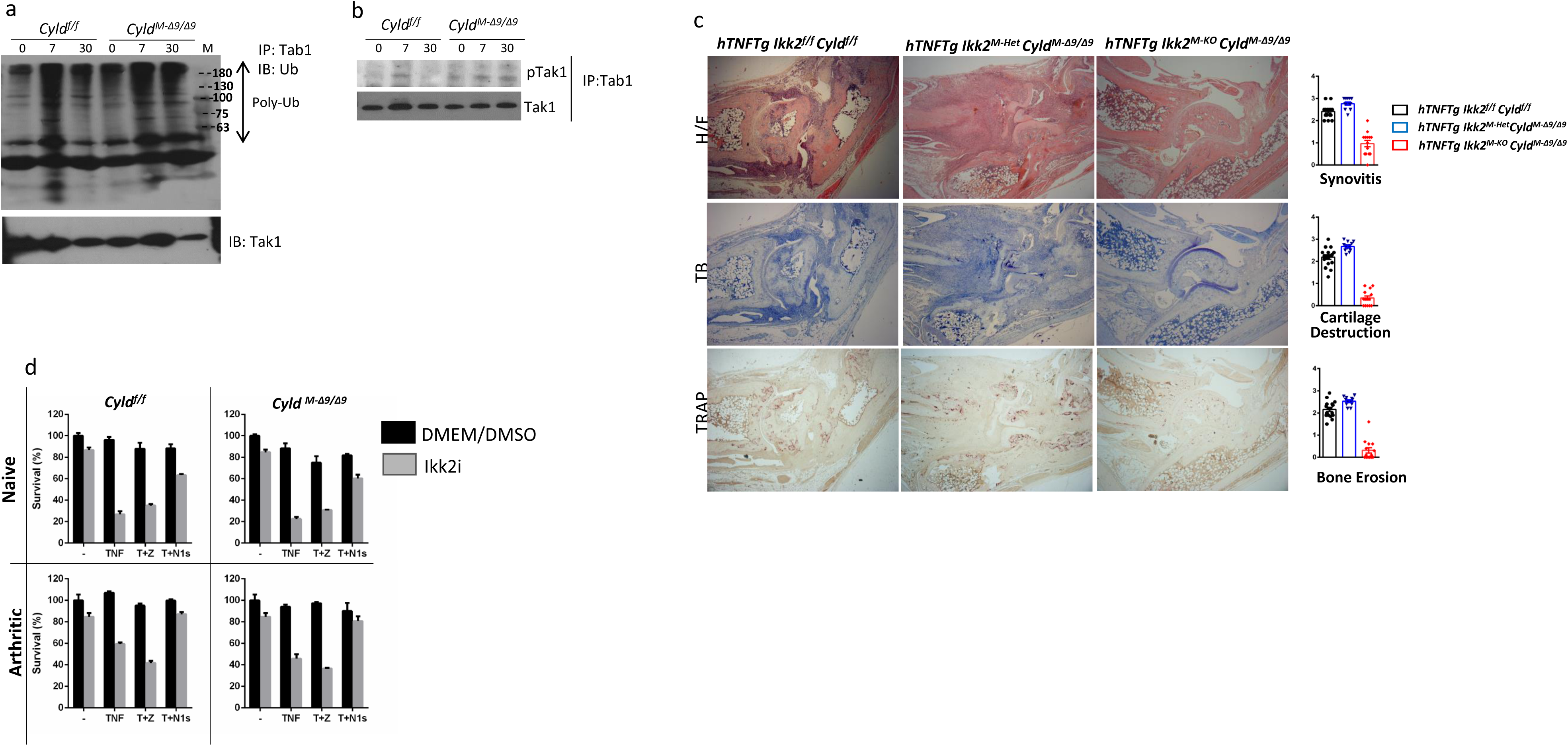
The TAK1/ikk2 axis regulate TNF arthritogenicity in TNF-mediated arthritis (a) Detection of Tak1 ubiquitination state upon TNF stimulation in lysates from *Cyld^f/f^* and *CyldM-Δ^9/Δ9^* SF cultures (subjected to IP with Tab1 antibody), for the indicated time points. (n=3) (b) Detection of phosphorylation state of Tak1 upon TNF stimulation in lysates from *Cyld^f/f^* and *Cyld^M-Δ9/Δ9^* SF cultures (subjected to IP with Tab1 antibody), for the indicated time points. (c) Histological evaluation *of hTNFtg Ikk2^f/f^ Cyld^f/f,^ hTNFtg Ikk2^M-Het^Cyld^Μ-Δ9/Δ9^* and *hTNFtg Ikk2/Cyld^Μ-Δ9/Δ9^* mice (d) Survival rates of *Cyld^f/f^* and *Cyld^M-Δ9/Δ9^* SF SFs treated with 50ng/ml TNF, in the absence (Black bars) of absence (Grey bars) of ikk2 inhibitor ML120b, zVAD and Nec1s as indicated (T+Z: TNF 50ng/ml + zVAD, T+N1s: TNF 50ng/ml+ Nec1s)(*n* = 3-5)

Since evidence in *hTNFtg* mice regarding either Jnk1^33^ or Jnk2 inactivation (Suppl Fig. 1) argues against a sufficient role of each JNK kinase activation for the development hTNF-mediated disease, we reasoned that the prolonged ubiquitination of TAK1 due to Cyld deficiency in SFs, causes the highly active state of IKK2 (Fig 4a, c), leading to enhanced NFkB and arthritogenic responses *in vivo*. To validate our speculation, we generated the mesenchymal-specific Cyld/Ikk2 mutants in the *hTNFTg* background. The arthritic phenotype of the *hTNFTg Ikk2* ^M-KO^*/Cyld*^M-Δ9/Δ9^ mice was modified and attenuated compared to their littermate Ikk2/Cyld-sufficient controls (Fig 5c), indicating that dominant Ikk2-mediated signaling is responsible for the exaggerated arthritic response in *hTNFTg Cyld*^M-Δ/Δ^ mice.

### Cyld is insufficient to block TNF-induced death of SFs

Previous evidence demonstrated that Cyld is implicated in the regulation of TNF-mediated programmed necrosis through the deubiquitination of Ripk1, abrogating the formation of the Ripk1/Ripk3 complex which activates the necroptosis executor kinase Mlkl^27^. In the context of hTNFtg arthritis, complete absence of Ripk3 kinase does not affect development of arthritis, indicating that Ripk3-mediated signals are not implicated in the pathology. However, Ripk3 can affect arthritic disease by promoting SF death only in the absence of IKK2 kinase^2^. Indeed, the arthritic phenotype of *hTNFTg Ikk2*^M-KO^ mice is characterized by residual inflammation, which is abrogated by Mlkl deficiency, confirming the inflammatory activity caused by necroptosis of IKK2-null SFs in vivo (suppl fig 2). In sharp contrast, SF-Cyld DUB deficiency was not equally sufficient to abrogate residual disease of *hTNFTg Ikk2*^M-KO^ mice (Fig. 5c). Consistent with the *in vivo* phenotype of *hTNFtg* mesenchymal-specific *Cyld/Ikk2* mutants, TNF-induced cytotoxicity in Cyld proficient and deficient SFs exhibited similar survival outcomes either in the presence or absence of IKK2 activity and caspases. Addition of Nec1s (Ripk1 inhibition), however, was sufficient to rescue lethality by TNF and Ikk2 inhibition in SFs of all genotypes examined^2^ (Fig 5d). This finding indicates that Cyld targeting is insufficient to abrogate the TNF-induced, Ripk1 mediated cytotoxicity in IKK2-null SFs and, consequently, the necroptosis-mediated residual disease in the *hTNFtg* mesenchymal-specific Ikk2 mutants.

Collectively, our results show that SF-Cyld controls TAK1 kinase to prevent IKK2 and NFkB hyperactivation but not Ripk1-mediated death, leading to deterioration of arthritic phenotype in TNF-dependent arthritides.

## Discussion

It is now well acknowledged that the persistent inflammation in RA entails a positive feedback whereby activated SFs recruit and retain immune cells to the joint, causing both cell types to shape and respond to the microenvironment in a self–sustaining manner^34^. Therapy for RA still remains problematic, owing to the poor understanding of the underlying activation mechanisms of SFs. Even though studies on the role of Cyld-mediated deubiquitination revealed important contributions in signaling pathways involved in development, homeostasis and several pathologies^8^, less is known about the its role in inflammatory arthritis. CYLD is significantly downregulated in human cultured RASFs compared to Osteoarthritic SFs, and this correlates well with the enhanced NFkB-mediated responses of RASFs^35^. How this correlation of SF CYLD and NFkB is mechanistically decoded and what is the factual effect of SF CYLD DUB deficiency in *in vivo* settings are still missing. Here we provide evidence supporting a role for CYLD in SFs as a new and robust modulator of inflammatory arthritis that functions to coordinately limit excessive activation of inflammatory and destructive properties of SFs. We show that mesenchymal Cyld largely contributes to prevent excessive JNK and NFkB activation in TNF-stimulated naive and arthritic SFs. Accordingly, high transcriptional activity of proteases and cytokines involved in the arthritic process accompanied the deteriorated phenotype of mesenchymal-Cyld DUB deficient mice. These effects are observed both in genetic and induced models of arthritis. In sharp contrast, the elimination of Cyld DUB activity in the mesenchymal tissues (Col6a1Cre+) is not detrimental for homeostasis.

The molecular basis of Cyld DUB activity in limiting JNK and NFkB prolonged activation in SFs highlights a common denominator of the two pathways, the Tak1 kinase. Tak1 hyperactivation occurs only upon TNF treatment of SFs and mediates sustained IKK2 phosphorylation that ultimately leads to activation of NFkB. a transcription factor that predominantly carries the arthritic potential of SFs in TNF-mediated arthritis^2,4^. Even though JNK/MAPK pathway is deregulated in Cyld-proficient SFs, our results and previous studies argue against a dominant pathogenic role of either JNK2 or JNK1 in TNF-mediated arthritis^33^, without excluding, however, their synergistic role in disease. Moreover, other mechanisms, including reactive oxygen species (ROS) accumulation, may also contribute to increased JNK phosphorylation^36,37^. Consistent with our findings on TAK1 hyperactivation, previous pharmacological studies have identified TAK1 as a central signal transducer that mediates the pathogenic activation of SFs *ex vivo* and the arthritic outcome *in vivo*^38,39^. We failed to generate double genetic mesenchymal mutants of Cyld/Tak1 that would allow us to further analyze the mesenchymal Cyld/TAK1 crosstalk in arthritis, as previously exhibited for hepatocytes^26,28^, due to embryonic lethality and the absence of inducible SF-specific Cre lines.

In contrast to the embryonic lethality of mesenchymal Cyld/Tak1 mutants, the concomitant Cyld/Ikk2 mesenchymal deficiency indicated the cause of exaggerated arthritic responses under mesenchymal Cyld-DUB deficiency. This approach also suggested that Cyld DUB deficiency was not sufficient to block the de novo RIPK3/MLKL-dependent necroptosis detected in Ikk2-targeted arthritic SFs *in vivo* and *ex vivo*^2^, contrasting current concepts on a dominant role of Cyld in regulating necroptosis. However, several studies support the context-specific effects of complete Cyld protein deficiency in different cell types, such as gingival fibroblasts in periodontal diseases, where TNF and LPS exerted differential NFkB outcomes^40^ or in Cyld-deficient macrophages and T-cells which differentially responds to innate (TNF, LPS) or adaptive (anti-CD3) stimuli respectively^41,42^. With regards to the regulation of death responses, the RIPK3-mediated death observed in the colon of intestinal epithelial NEMO deficiency was not rescued by Cyld DUB deficiency^43^, while the repression of CYLD in neuronal cells promoted cell death but did not alter NF-κB activity upon brain injury^44^. However, we cannot fully dismiss that incomplete Cyld targeting, dominant-negative functions of truncated CYLD^28^, other spatiotemporal signaling events^45^ or context-specificity of signaling counterparts of Cyld (Itch1, usp18, USP4, spata2) account for the insufficiency of Cyld to prohibit necroptosis in our experiments with the Ikk2-deficient SFs.

In the same line of evidence, It is also tempting to consider that K63 deubiquinating activity of A20 cannot reconstitute for the Cyld insufficiency in SFs, indicating that the crucial target of Cyld in SFs could be other than the common set of targets such as TRAF2, TRAF6, RIP1, and NEMO^8^. It is previously suggested that A20 and Cyld differentially acts to regulate TNF pathway; A20 expression is induced by inflammatory stimuli and acts to terminate responses^46,47^ while Cyld is constitutively active to prohibit uncontrolled NFkB activation^48^. Alternatively, a prominent role of Cyld-mediated M1 deubiquitination in TNF signalling of SFs remains to be resolved^49,50^.

Collectively, by exhibiting the major contribution of Cyld deubiquitinase activity in the TAK1/IKK2 axis leading to pathogenic NFkB, we identify Cyld as a negative regulator of the major inflammatory signaling acting downstream of Tnfr1 in SFs. Our study, therefore, contributes to the signaling map underlying SF responses in TNF-mediated arthritis that would inform potential therapeutic intervention strategies targeting SFs.

## Material and Methods

### Animals and Induction of Arthritis

Human TNF transgenic (*hTNFtg*)^31^, *ColVI-Cre*^3^, Cyld^flx9/flx9^, Milk^−/−^^51^ and Ikbkb ^f/f^^52^ (referred in the text and figures as Ikk2^f/f^) mice have been previously described. *ColVI-Cre* Cyld^flx9/flx9^ mice referred to as Cyld^M-Δ9/Δ9^. Collagen-Antibody-Induced-Arthritis (CAIA) was induced according to manufacture instructions (MD Biosciences). Mice were maintained on a C57BL/6J genetic background.

### Clinical Assessment of Arthritis

Arthritis in *hTNFtg* animals was evaluated in ankle joints in a blinded manner using a semiquantitative arthritis score ranging from 0 to 4; 0: no arthritis (normal appearance and grip strength); 1: mild arthritis (joint swelling); 2: moderate joint swelling and digit deformation, reduced grip strength; 3: severe joint swelling and digit deformation, impaired movement, no grip strength; 4: severe joint swelling and digit deformation, impaired movement, no grip strength and obvious weight loss-cachexia At indicated weeks of age, mice were killed and the hind ankle joints were removed for histology.

CAIA clinical evaluation was performed as previously described ^53^.

### Histologic Analysis

Formalin-fixed, EDTA-decalcified, paraffin-embedded mouse tissue specimens were sectioned and stained with hematoxylin and eosin (H&E), Toluidine Blue (TB) or TRAP kit [Sigma-Aldrich]. H&E-stained joint sections were semi-quantitatively blindly evaluated for the following parameters: synovial inflammation/ hyperplasia (scale of 0–4), cartilage erosion (scale of 0–4), and bone loss (scale of 0–4)^54^.

### Microcomputed tomography

Microcomputed tomography (mCT) of excised joints was carried out by a SkyScan 1172 CT scanner (Bruker, Aartselaar, Belgium) following the general guidelines used for assessment of bone microarchitecture in rodents using mCT ^55^. Briefly, scanning was conducted at 50 kV, 100 mA using a 0.5-mm aluminum filter, at a resolution of 6 μm/pixel. Reconstruction of sections was achieved using the NRECON software (Bruker) with beam hardening correction set to 40%. The analysis was performed on a spherical volume of interest (VOI) (diameter 0.53mm) within 62 slides of the trabecular region of calcaneus. Morphometric quantification of trabecular bone indices such as trabecular bone volume fraction (BV/TV %), trabecular number (Tb. N; 1/mm) and trabecular separation (Tb. Sp; mm) were performed using the CT analyzer program (Bruker).

### Flow cytometric analysis of joint tissue

The ankle joint tissues were disaggregated by incubation for 40 min at 37°C in an enzymatic digestion medium consisting of DMEM, 10%heat-inactivated FBS, collagenase type IV (300 units) from *Clostridium histolyticum* [Sigma] and 0.03 mg ml^-1^ DNase [Sigma]. Upon washing with PBS containing DNase, the cells were blocked in 1% BSA in PBS and Fc blocker (anti-CD16/32, Biolegend 101302) for 10 min at 4°C. For SF subtype determination, the following fluorophore conjugated antibodies were employed: anti-Pdpn PE-Cy7, 127411; anti-Thy1 A647, 105318; anti-CD31 APC/Fire 750, 102433; anti-CD45 APC-Cy7, 103116 [Biolegend]; anti-Ter119 APC-A780, 47-5921-80 [eBioscience]. For myeloid infiltrates, we used the following antibodies [Biolegend]: anti-Ly-6C FITC, 128005; anti-NK-1.1 PE, 108707; anti-CD11c PE/Dazzle™ 594, 117347; anti-Ly-6G PE/Cyanine7, 127617; anti-CD11b APC, 101211; anti-CD45 APC/Cyanine7, 103116. The analysis was performed with BD FACSCanto II and the BD FACSDiva software and dead cells were excluded by Propidium Iodide staining. Analysis of the results were performed employing FlowJo software (v.10)

### Cell culture

Primary mouse SFs were isolated from mice with indicated genotypes and cultured for four passages as previously described^56^. Upon evaluation of purity (less of 5% CD45+ cells in the culture), SFs were used in the assays.

In experiments with Ikk2 inhibitor, pretreatment of cells with ML120B (Tocris, 10uM)^57^ was performed for 3h before addition of other reagents. zVAD [Bachem] and Nec1s [BioVision] were used in concentrations 20 and 30mM respectively. TNF cytotoxicity assays were performed in 96-well plates, terminated 16-18h post TNF addition, and evaluated upon crystal violet staining, solubilization in 33% acetic acid solution and determination of optical density at 570nm as a test filter and 630nm as a reference filter.

Flow cytometric analysis was performed in cells stained with the following antibodies CD90.2, VCAM-1 [eBioscience], CD45 [BioLegend], ICAM-1 and CD45 [BD Biosciences] using FlowJo analysis software.

### Immunoblotting and Immunoprecipitation

For immunoblotting, samples were collected in RIPA buffer, containing 1% Triton X-100, 0.1% SDS, 150 mM NaCl, 10 mM Tris HCl, pH 7.4, 1 mM EDTA, protease inhibitors [Roche], and phosphatase inhibitors [Sigma-Aldrich], separated by SDS/PAGE (10-12.5%), transferred to nitrocellulose membranes [Millipore], and probed with the following antibodies: Ikk2, pJNK1/2, p-Ikk1/2, p-p38 [Cell Signaling]; ikB, JNK, p38, p-ERK1/2, ERK1/2, tubulin, actin [Santa Cruz Biotechnology].

For immunoprecipitation experiments, SFs were stimulated with murine TNF (20ng/ml) (VIB Protein Service Facility) as indicated. Cells were lysed in NP40 buffer (150 mM NaCl, 1% NP40, 10% glycerol and 10 mM Tris–HCl pH 8), and the extracts were incubated with 2ug anti-Tab1 antibody^58^ (kindly provided by Philip Cohen/University of Dundee) for 3h. Protein A/G beads [Santa Cruz Biotechnology] were added for the 5.5 h of incubation. The beads were then washed three times and the samples were eluted from the beads using Laemni buffer, separated by SDS/PAGE (8%), transferred to nitrocellulose membranes and subjected to immunoblot, as described, utilizing antibodies against Tak1, p-Tak1 [Cell Signaling] and ubiquitin [Santa Cruz Biotechnology].

### EMSA

To generate dsDNA with NFkB binding site, we mixed appropriate oligos with complementary sequences (200ng each) (sense 5’-ATCAGGGACTTTCCGCTGGGGACTTT-3’ and antisense 5’CGGAAAGTCCCCAGCGGAAAGTCCCT-3’), heated to 100 °C, and allowed to return slowly to room temperature. Upon ^32^P-labeleling, the ds DNA fragments with the NFkB binding site were incubated with nuclear extract (5 μg per reaction) in 25 mM Hepes-KOH at pH 7.9, 250 mM KCl, 25 mM MgCl_2_, 2.5 mM DTT, 50% glycerol, at room temperature for 20 min. After addition of 20% ficoll, samples were subjected to electrophoresis on a 6% polyacrylamide gels run with 50 mM Tris-borate buffer, pH 8.3, 1 mM EDTA at 4°C. Dried gels were visualized by phosphoimaging using Personal Molecular Imager FX system (Bio-Rad Laboratories) or films.

### Statistical analysis

Data are presented as mean ± SE. Analyses were performed by employing SigmaPlot and GraphPad software. P-values <0.05 were considered significant.

### Study Approval

Experiments were performed in the conventional unit of the animal facilities in Biomedical Sciences Research Center (BSRC) “Alexander Fleming” under specific pathogen–free conditions, in accordance with the guidance of the Institutional Animal Care and Use Committee of BSRC “Alexander Fleming” and in conjunction with the Veterinary Service Management of the Hellenic Republic Prefecture of Attika/Greece (License Νο: 4405-5.7.2016). Experiments were monitored and reviewed throughout its duration by the respective Animal Welfare Body for compliance with the permission granted.

## Acknowledgements

The authors thank Panos Athanasakis, Spyros Lalos and Anna Katevani for excellent technical assistance. Authors also thank George Kollias for critical discussions and for providing reagents and the hTNFtg model, Manolis Pasparakis (University of Cologne, Germany) for kindly providing the Ikk2^fl/fl^ and Mlkl^−/−^ mice, Claude Libert (VIB, Ghent, Belgium) for the recombinant TNF and Dimitris Kontoyiannis (Aristotle University of Thessaloniki, Greece) for critical reading of the manuscript. This work has been supported by the grant from the Stavros Niarchos Foundation to the Biomedical Sciences Research Center “Alexander Fleming” (startup fund to M.A.), as part of the Foundation’s initiative to support the Greek research center ecosystem, and FOREUM grant (StroPHe, #061 to MA). The authors also wish to acknowledge the support from the InfrafrontierGR infrastructure (co-financed by the ERDF and NSRF 2007-2013) for providing excellent mouse hosting, transgenesis, flow cytometry, histology and microCT facilities.

## Figure Legends

Suppl. Figure 1. Jnk2 does not critically regulate the arthritic phenotype of *hTNFtg* mouse Histological evaluation of *hTNFtg Jnk2*^+/−^ and *hTNFtg Jnk2^−/−^* mice with H/E and toluidine blue stainings (n=5-6) at age of 8 weeks.

Suppl. Figure 2. Mlkl is necessary to prevent residual disease in *hTNFtg* Ikk2^M-KO^ mice. Histological evaluation of *hTNFtg Mlkl−/+*, *hTNFtg Mlkl−/−, hTNFtg Mlkl^+/+^ Ikk2^Μ-Δ9/Δ9^*and *hTNFtg Mlkl^−/−^* Ikk2^Μ-ΚΟ^ with H/E and toluidine blue stainings (n=6-12) at age of 10 weeks.

