## Supplementary figures and images for "Cyld aborts hyperactivation of synovial fibroblasts in inflammatory arthritis by regulating the TAK1/IKK2 signaling axis"

### Supplemental Figures

Suppl Figure 1

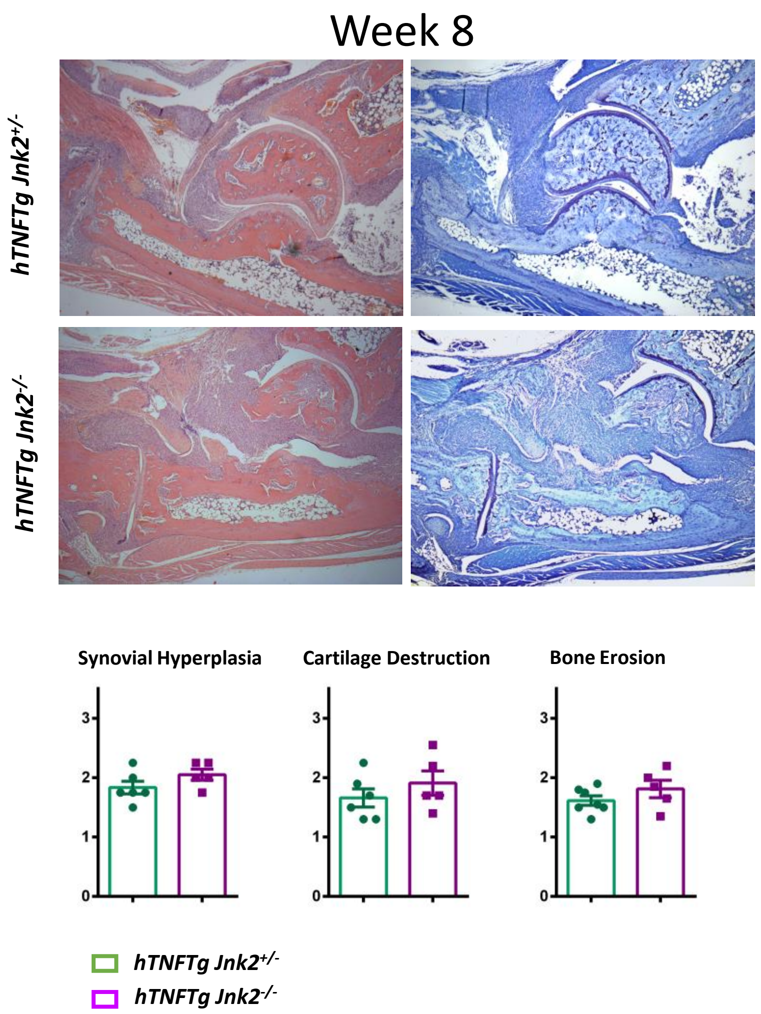

#### Suppl Figure 2

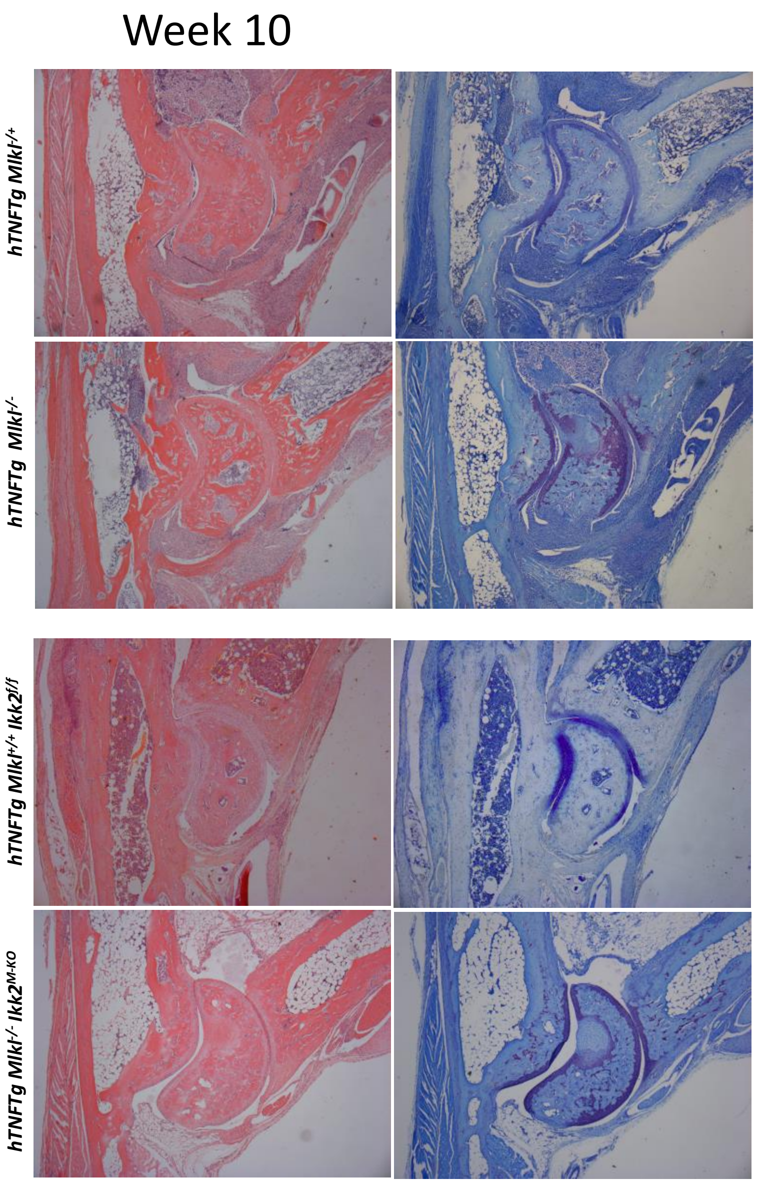


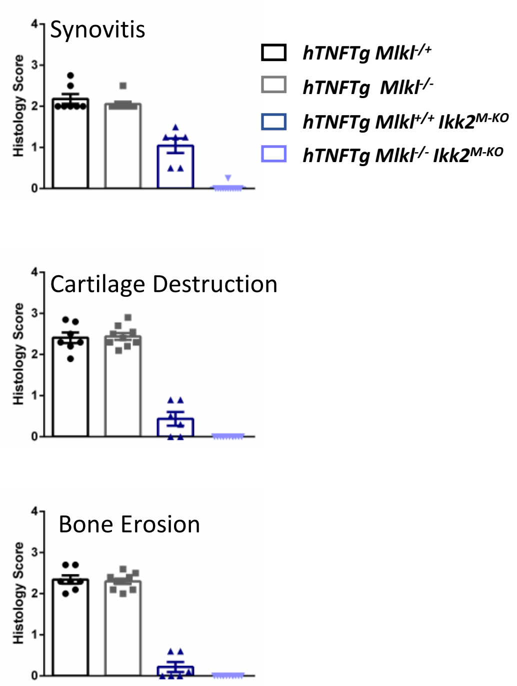
